# PolyA tracks and poly-lysine repeats are the Achilles heel of *Plasmodium falciparum*

**DOI:** 10.1101/420109

**Authors:** Slavica Pavlovic Djuranovic, Jessey Erath, Ryan J Andrews, Peter O Bayguinov, Joyce J Chung, Douglas L Chalker, James AJ Fitzpatrick, Walter N Moss, Pawel Szczesny, Sergej Djuranovic

**Author notes:** equally contributed (Slavica Pavlovic Djuranovic, Jessey Erath).

## Abstract

*Plasmodium falciparum*, the causative agent of human malaria, is an apicomplexan parasite with a complex, multi-host life cycle. Sixty percent of transcripts from its extreme AT-rich (81%) genome possess coding polyadenosine (polyA) runs, distinguishing the parasite from its hosts and other sequenced organisms. Recent studies indicate that transcripts with polyA runs encoding poly-lysine are hot spots for ribosome stalling and frameshifting, eliciting mRNA surveillance pathways and attenuating protein synthesis in the majority of prokaryotic and eukaryotic organisms. Here, we show that the *P. falciparum* translational machinery is paradigm-breaking. Using bioinformatic and biochemical approaches, we demonstrate that both endogenous genes and reporter sequences containing long polyA runs are efficiently and accurately transcribed and translated in *P. falciparum* cells. Translation of polyA tracks in the parasite does not elicit any response from mRNA surveillance pathways usually seen in host human cells or organisms with similar AT content. The translation efficiency and accuracy of the parasite protein synthesis machinery reveals a unique role of ribosomes in the evolution and adaptation of *P. falciparum* to an AU-rich transcriptome and polybasic amino sequences. Finally, we show that the ability of *P. falciparum* to synthesize long poly-lysine repeats has given this parasite a unique protein exportome and an advantage in infectivity that can be suppressed by addition of exogenous poly-basic polymers.

## Main

The complex life cycle of *Plasmodium falciparum*, responsible for 90% of all malaria-associated deaths, involves multiple stages in both the human and mosquito hosts. Asexual replication during the intraerythrocytic development cycle (IDC) is tightly regulated over a 48-hour period; involving the expression of the majority of its genes. Progression through asexual stages (ring, trophozoite, schizont) of the IDC requires a strictly controlled panel of gene expression profiles for each stage. A range of 16-32 daughter cells results from the IDC. Thus, a single, originating merozoite must undergo several rounds of DNA synthesis, mitosis, and following nuclear division in a relatively short period (*1-3*). The apparent necessity for rapid and accurate DNA replication, RNA transcription, and protein translation, as well as competent folding machinery, is a further emphasized by the recent demonstration that genes important for these processes are essential in *P. falciparum* (*4*). Even so, faithful execution of these fundamental processes is challenged by the AT-rich *P. falciparum* genome: averaging ∼81% overall AT content: ∼90% in the non-coding region and, a still relatively high, ∼76% within the coding region (*5-7*). Recently, it was demonstrated that the translation of genes with polyadenosine runs (polyA tracks), primarily coding for lysine residues, is attenuated in the majority of tested eukaryotic and prokaryotic organisms due to ribosomal stalling and frameshifting on such RNA motifs (*8-15*). In humans, the presence of just 12 adenosines in an mRNA coding region will deplete the protein synthesis of a given gene by more than 40% (*9, 10*). The consequence of translational arrest is activation of one or more mRNA surveillance mechanisms and has been demonstrated in studies of human, yeast and *E. coli* (*9, 12*).

High AT-content within coding regions and an extreme AAA and AAT codon bias, increases the propensity for polyA tracks in the *P. falciparum* transcriptome (*16*). Additionally, a “just-in-time” transcriptional model of gene expression in *P. falciparum* has been proposed whereby a transcriptional burst produces most of the gene transcripts required for the IDC during the trophozoite stage (*2, 17*). While both the DNA and RNA polymerases must contend with high DNA AT-content, it is even more interesting how the unusual AU-richness of *Plasmodium* mRNAs would impact the fidelity and efficacy of protein synthesis. With “just-in-time” translation of numerous A-rich coding sequences (*7*) and poly-lysine proteins harboring an AAA codon bias, expressed at all stages in the parasite life cycle (*2, 3, 18*), the *Plasmodium* translation machinery represents a paradigm-breaking system in protein synthesis compared to other organisms. Given the high impact of possible changes on both nucleic acid and protein metabolism, driven by enrichment of polyA and poly-lysine sequences, *Plasmodium* species must have adapted their protein synthesis machinery to this trait. Furthermore, such evolutionary inherited features in the *P. falciparum* parasite must have resulted in survival and reproduction benefits for this organism in its particular environment(s).

In this study, we aimed to determine translational efficiency and accuracy of protein synthesis from polyA tracks in *P. falciparum* cells and explored differences in the translational machinery that may be evolved to accommodate the unusual AU-richness of the *P. falciparum* transcriptome. Furthermore, we sought to delineate benefits of poly-lysine repeats, resulting from the translation of polyA tracks, on parasite RNA metabolism, survival, and pathogenicity.

## Results

While the underlying reasons for the disproportionate representation of the four nucleotides in any given genome may be different, it is of vital importance that shifts towards extreme AT- or GC-richness must be accommodated by adaptation of the transcription and translation apparatuses; enabling the cell to transcribe and translate each gene appropriately. Characterization of gene organization in *P. falciparum* revealed that both coding and non-coding regions contribute equally to overall AT-richness of its genome (*7*). To explore the association between polyA tracks—focusing on ≥12 adenosine nucleotides in coding sequences that have previously been shown to induce stalling, as well as genomic AT-content—we analyzed 250 eukaryotic genomes (Fig. 1A). As we hypothesized, *P. falciparum* and genus affiliates have a much higher ratio of polyA track genes when normalized to genomic AT-content. This feature of *Plasmodium* species is conserved regardless of their genomic AT-content, resulting in two groups (low and high AT-content *Plasmodium spp*.) with unusually high portions of polyA track genes ranging from 35 to over 60% of the total transcriptome. To further emphasize the differences between *P. falciparum* and other organisms, we have analyzed genes that contain polyA tracks and their full length (Supplementary Fig. 1 and Fig. 1B, respectively).

**Figure 1.**
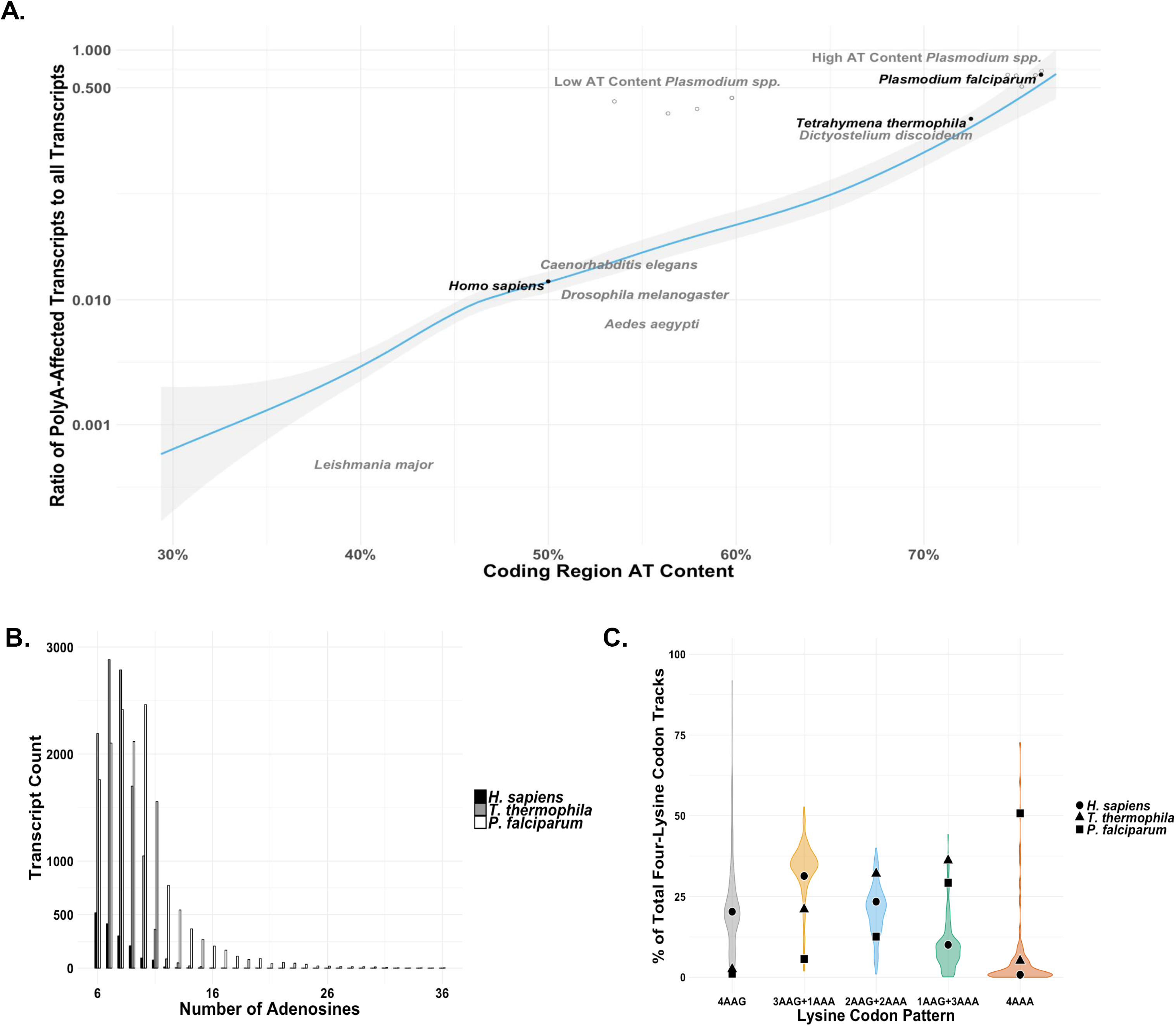
**A.** The plot of 152 species representing a comparison of the ratio of polyA-affected transcripts (over a total number of transcripts) to the AT content of coding region for each organism. Organisms pertinent to this paper are in black. For reference, other model organisms of interest are displayed in gray, including a position of various *Plasmodium spp*. **B.** Transcript counts for genes with 6 to 36 consecutive adenosines for *H. sapiens, T. thermophila,* and *P. falciparum. H. sapiens and T. thermophila* are limited to a single transcript at length of ≤17As. The longest P. falciparum transcript reaches maximal 65As, with multiple transcripts of ≤36As. D. Violin plot of lysine codon usage distribution in tracks of four lysine residues for 152 organisms. *H. sapiens* (circle), *T. thermophila* (triangle), and *P. falciparum* (square) are specifically noted.

In comparison with the human host and *Tetrahymena thermophila*, another protozoan with an AT-rich genome, *P. falciparum* showed an immense amount of polyA track genes (12 adenosine nucleotides allowing for one mismatch - 12A-1, Supplementary Fig. 1). Around 60% of genes in *P. falciparum* have ≥12 A’s in a row, while humans and *T. thermophila* range from just 2-5%. This difference is even more apparent in transcript counts that host polyA tracks (Fig. 1B). The *P. falciparum* genome contains more than 1000 genes with 16 consecutive adenosine nucleotides (16As); reaching a maximum of 111 As in *P. falciparum fch 4* strain (*19*) (Fig. 1C).

The observed disparity in the number of genes with polyA tracks could be due to previously observed codon bias in *P. falciparum* (*16*). However, it was already shown that codon bias and tRNA abundance do not correlate with codon selection in genes coding for lysine repeats (*9, 12*). To investigate in more detail the distribution of AAA and AAG codons in polylysine tracts, we have analyzed transcripts from *P. falciparum* and other eukaryotic genomes (Fig. 1C). We observed a complete reversal of the trend exhibited in other organisms, including humans and *T. thermophila*, with the highest abundance of *P. falciparum* transcripts hosting four consecutive AAA codons in stretches of four lysine residues (Fig. 1C). This divergence from other analyzed transcriptomes is preserved in other members of *Plasmodium spp*., with *P. berghei* being an extreme example using only AAA codons in 73% of transcripts coding for poly-lysine stretches. A majority of *Plasmodium* poly-lysine proteins fall into the group of essential genes based on the recent mutagenesis studies (*4*). This outcome is expected given that gene ontology results indicate enrichment in gene products involved in the crucial cellular process such as translation initiation, chromosome segregation, previously observed in other organisms (*9*) as well as in cellular and pathological cell adhesion. Interestingly, cellular and pathological adhesion genes that are the hallmark of *Plasmodium* infectivity and pathogenicity came as the only enriched biological process in gene ontology search with polyA track carrying genes (Supplementary Table 1). Previous analysis indicated that 70%-85% of orthologs of polyA carrying genes from *P. falciparum* have the same polyA segments in genes from other *Plasmodium* species, regardless of their genomic AT content (*19*). This high level of conservation is not found in other organisms arguing for the possible functional role of polyA tracks and poly-lysine repeats within *Plasmodium* species. As such, our bioinformatic analyses demonstrate that conservation of both polyA tracks in the transcriptome and poly-lysine repeats in the proteome of *P. falciparum* have been evolutionary selected and conserved due to the possible benefits for the parasite.

Given the diversity of RNA functions, mRNA stability and successful translation are made possible, and affected, by the ability of RNA to fold into unique functional structures (*20*). Shifts towards AU-/GC-richness may affect the propensity and stability of RNA structures thermodynamically. This possible effect on RNA begs the question as to whether or not the presence of polyA tracks, and thereby poly-lysine repeats, provide important effects on mRNA structure and, subsequently, protein synthesis in *P. falciparum*. To examine how high AU-content influences predicted mRNA structure and stability in different organisms, we compared the ability of all mRNAs to form thermodynamically stable structures and calculated their stability using *in silico* approaches. The coding transcripts from *T. thermophila* and *P. falciparum* had on average higher (less stable) predicted minimum free energy (MFE, Figure 2A) of folding than transcripts from human; possibly due to the AU-bias of these organisms. Interestingly transcriptome z-scores—measuring the difference in stability of native vs. random sequence—from the same organisms revealed no significant differences between mRNAs from each (z-score, Fig. 2B). This result indicates that while folding stability varies greatly between species (following from the skews in nucleotide content), the propensity of native sequences to be ordered to fold, does not.

**Figure 2.**
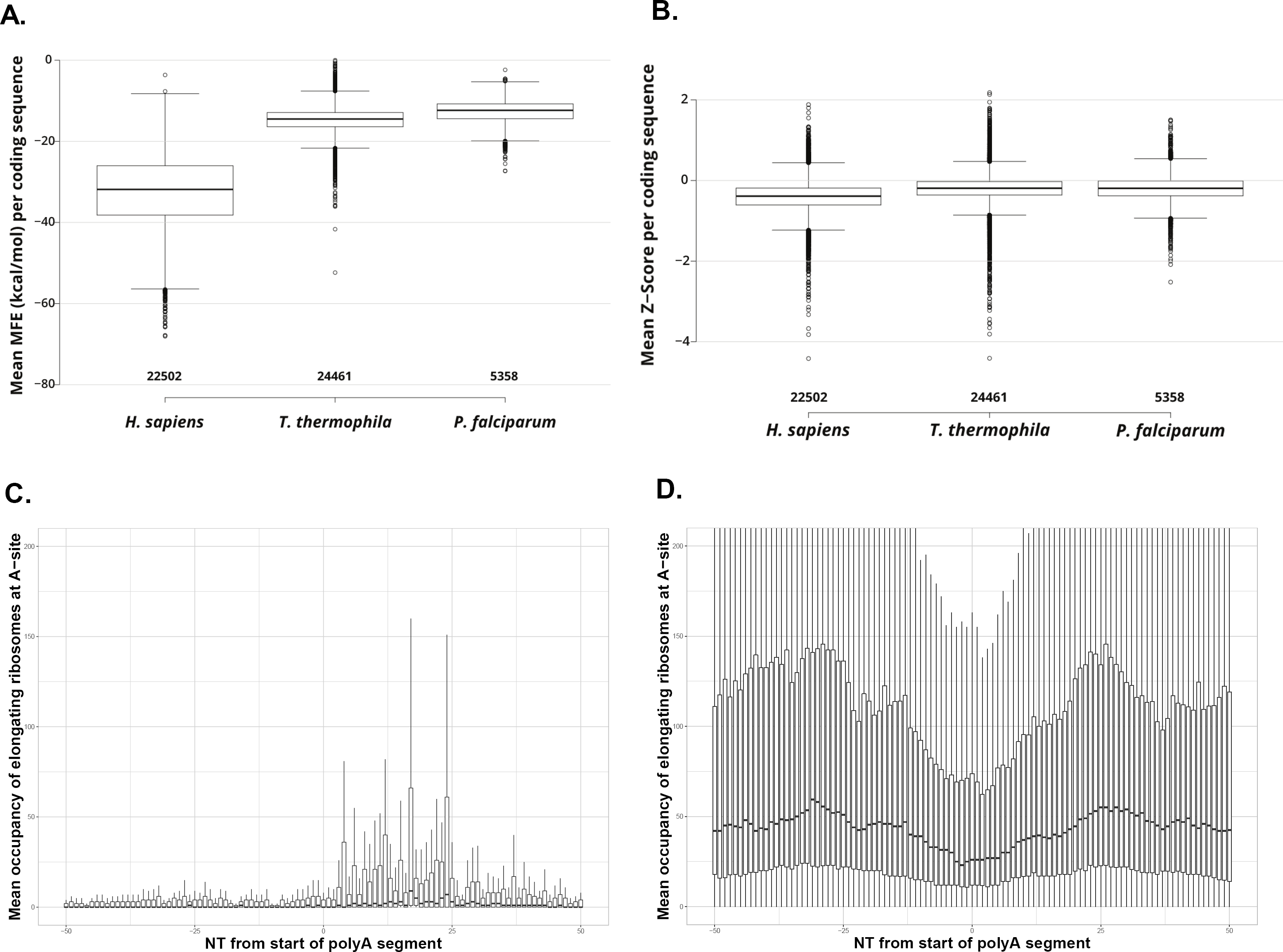
**A.** Box and whisker plots are showing the distribution of mean folding energy values (kcal/mol; measures the stability of RNA structure) calculated for each coding sequence from *H. sapiens, T. thermophile,* and *P. falciparum*, resulting from a scanning window analysis (see Methods). Center lines show the medians; box limits indicate the 25th and 75th percentiles; whiskers extend 1.5 times the interquartile range from the 25th and 75th percentiles; outliers are represented by dots; numbers below plots are the number of coding sequences for each species. **B.** Box and whisker plots showing the distribution of mean z-score values (measures standard deviations more stable native sequence RNA structure is vs. random) calculated for each coding sequence from *H. sapiens*, *T. thermophila*, and *P. falciparum*, resulting from a scanning window analysis (see Methods). Plots are structured as in (**A**). Occupancy of elongating ribosomes (mapped to A-site) around the start of polyA segments in human (**C**) and *P. falciparum* (**D**) – in the same scale. In both cases, to avoid inclusion of sparsely mapped segments, regions with average occupancy below the mean for the whole dataset were excluded. In the case of *P. falciparum* gene segments with polyA tracks shorter than 22 nucleotides were taken into account.

With as much as 60% of the parasite transcriptome harboring hot-spots of translational stalling, we performed a comparative analysis of ribosome profiling data from *P. falciparum* (*3*) and aggregated data for human tissues conveniently harmonized at GWIPS database (*21*) to determine whether endogenous polyA tracks and poly-lysine sequences, induce translational pausing. Ribosome stalling can be observed in the ribosome profiling data as an increase in the abundance of ribosome footprints on sequences that cause ribosomes to pause during translation (*22*). Cumulative data for all transcripts with polyA tracks from human cells indicates substantial translational pausing on these sequences (Fig. 2C). However, no evidence for ribosome stalling could be observed for *P. falciparum* transcripts accommodating polyA tracks with a length of ≤ 22 consecutive adenosine nucleotides (Fig. 2D). This translation phenomenon is independent of different stages of *P. falciparum* intraerythrocytic development (Supplementary Fig. 3) and ostensibly irrespective of polyA track, or poly-lysine, length to such a degree that cumulative transcript analysis becomes hindered by low sequence complexity of this region or reduced number of reads for the long polyA tracks (Supplementary Fig. 4). While we observe a relatively small increase in the number of elongating ribosomes on polyA segments in the late trophozoite and schizont stages of IDC (Supplementary Fig. 5), it’s unclear if these translate into larger protein expression.

Given the potential for negative selection against polyA tracks (*23*)(Guler JL, 2013), particularly in laboratory conditions, we selected a small subset of varied length, differentially expressed polyA track-containing genes for expression analysis (GAPDH - PF3D7_1462800 (control), 610 - PF3D7_1464200 (longest polyA track length 20 As), CRK5 - PF3D7_0615500 (longest polyA track length 20As of three present tracks), IWS1L - PF3D7_1108000 (longest polyA track length 31As of two present tracks). Using qRT-PCR analysis of total RNA, we analyzed the expression profiles of select endogenous genes in our *P. falciparum* Dd2 lab strain. A time course study of synchronized parasite culture indicated that the selected polyA track transcripts are efficiently transcribed at all time points when compared to the control gene (Supplementary Fig. 6). Additionally, Sanger sequencing and gel analysis of cloned cDNAs on the selected subset of endogenous *P. falciparum* genes revealed that the annotated polyA tracks remained intact in our lab strain.

To investigate further the *P. falciparum* transcriptome feature that distinguishes it most from other organisms, we used double HA-tagged reporter constructs with a 36 adenosine nucleotide (36As) insertion, coding for 12 lysine residues, between the tag and a fluorescent protein (Supplementary Fig. 7). As a control, we used only HA-tagged reporter proteins. Episomal expression of the reporter in all three organisms was followed by analysis of RNA abundance by qRT-PCR and protein abundance by western blot detection of the HA-tag (Figure 3A and Figure 3B, respectively). As expected, we observed robust changes in mRNA levels (Figure 3A) and substantial losses in protein expression (Fig. 3B) for reporters with polyA tracks in both neonatal human dermal fibroblasts (HDFs) and AT-genome rich *T. thermophila* (*10, 11*). We observed minimal, if any effects, of polyA track insertion on reporter mRNA and protein expression in *P. falciparum* (Fig. 3A and 3B). Further analysis of *P. falciparum* cells by live-fluorescence microscopy confirms equivalent mCherry reporter expression, judging by the intensity of fluorescence, between constructs with and without polyA tracks (Fig. 3C). To assess whether the efficiency of polyA track translation is altered when located further downstream of the start codon, we designed a construct with thioredoxin and nano-luciferase (nanoluc) proteins separated with HA-tag and AAA-coded poly-lysine stretch (Supplementary Fig. 8). Measurement of nanoluc luminescence from the same number of drug-selected parasites indicates slightly higher expression of a reporter with polyA track compared with control reporter (Fig. 3D). The same ratio was observed in western blot analysis of protein, respectively (Fig. 3D and Supplementary Fig. 8). Taken together, our ribosome profiling analyses, selected endogenous gene expression analysis, and reporter expression data (Fig. 2 and 3) indicate that polyA tracks are tolerated by the parasite translational machinery, and suggest an adaptation of the *P. falciparum* translation and mRNA surveillance machinery to its AU-rich transcriptome and poly-lysine rich proteome.

**Figure 3.**
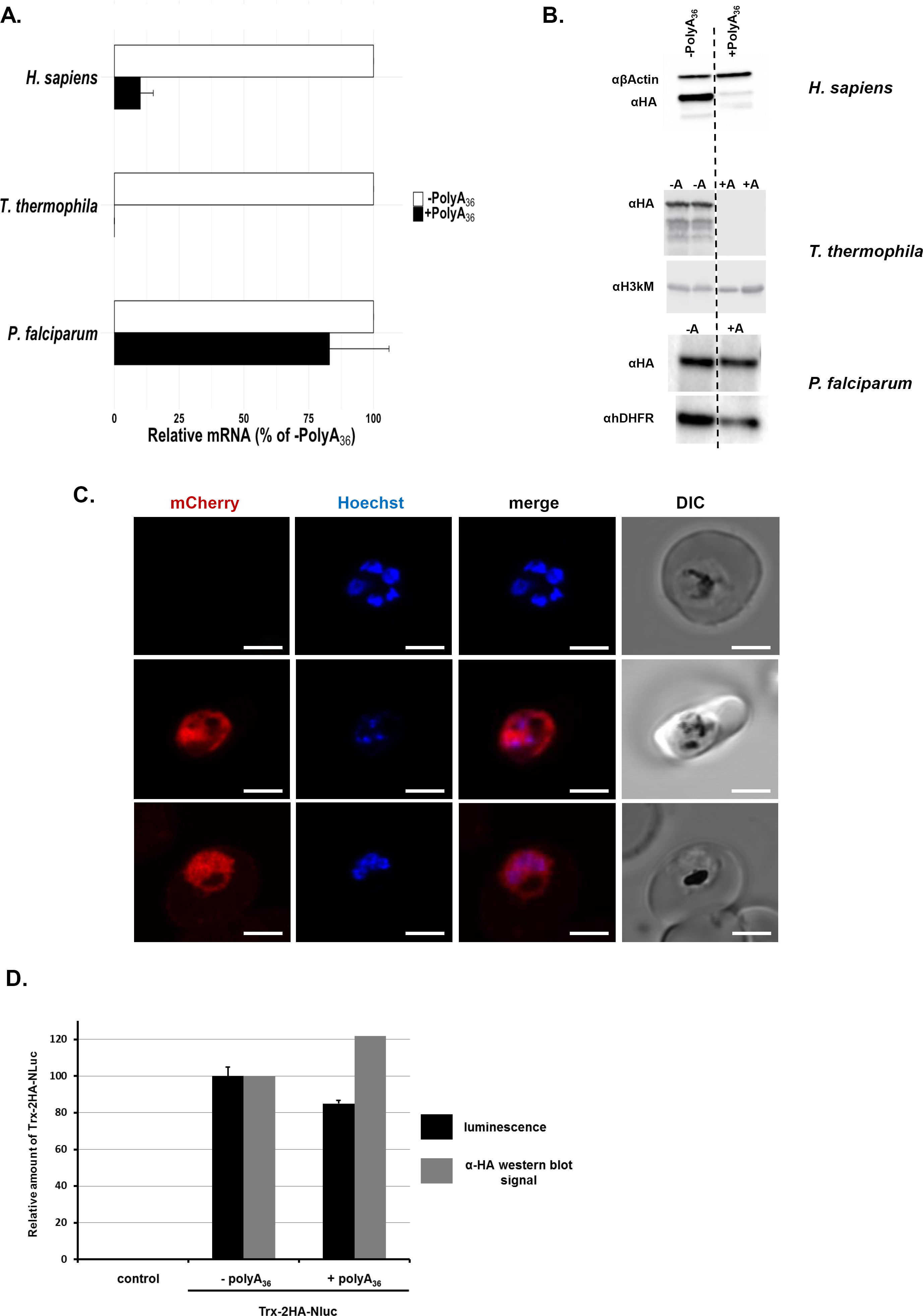
**A.** mRNA abundance of reporter constructs (+polyA36) by qRT-PCR relative to their counterpart lacking polyA stretches (-polyA36) in *H. sapiens, T. thermophila,* and *P. falciparum* cells. Data represent three biological replicates with a standard deviation. **B.** Expression of reporter constructs in *H. sapiens, T. thermophila,* and *P. falciparum* followed by western blot analysis with αHA or αGFP antisera. Samples from two integrated clones for the -polyA36 control (-A) and the +polyA36 reporter (+A) are shown for *T. thermophila*. αβ-actin, α-acetylated alpha-tubulin (ATU) and αhDHFR are used as loading controls for western blot analysis from *H. sapiens*, *T. thermophila* and *P. falciparum* cells, respectively. **C.** Images from live fluorescence microscopy of *P. falciparum* expression of reporter constructs with (+PolyA36) and without (-PolyA36) polyA tracks as well as parent (non-transfected) line, 2.5 µm scale bar. **D.** Quantification of protein amounts for Thioredoxin-2HA-NanoLuciferase (Trx-2HA-NLuc) reporter without (-polyA36) and with 36 adenosine stretch (+polyA36) expressed in *P. falciparum cells*. Western blot analysis of Trx-2HA-NLuc reporter (Supplementary Figure 11) and luminescence measurements were normalized to hDHFR or cell number, respectively. Luminescence data represents mean value of three biological replicates with standard deviation.

It has been shown the translation of polyA tracks elicits non-sense mediated decay (NMD) and “no-go” decay (NGD) mRNA surveillance pathways in other organisms (*8, 9, 12, 24, 25*). To test how *P. falciparum* has evolved and adapted to accommodate both polyA tracks and poly-lysine repeats, we first turned towards mRNA surveillance pathways and ribosomes as principal components that control mRNA quality, as well as the fidelity and efficiency of protein synthesis. While previous work demonstrated that the NMD pathway is intact (*26*), the existence of the NGD pathway was previously untested in *P. falciparum*. To explore whether changes in the NGD pathway are potential adaptations of *P. falciparum* to polyA tracks and poly-lysine repeats, we used CRISPR/Cas9 technology (*27, 28*) to HA-tag the endogenous Pelota homolog (PfPelo, PF3D7_0722100, (Supplementary Fig. 9 and 10) in *P. falciparum* Dd2 cells. The Pelo protein in complex with Hbs1, for which there is no clear homolog observed in *P. falciparum*, recognizes and rescues stalled ribosomes on long polyA stretches (*24, 25*). Increased Pelota recruitment to polysome fractions is one of the indicators of a stalled protein synthesis while the transcripts responsible for ribosomal stalling are targeted for the mRNA decay (*29, 30*). In the absence of sequences that directly stall *P. falciparum* ribosomes, isoleucine (Ile) starvation was used to assess the recruitment of PfPelo protein to stalled ribosomes (Supplementary Fig. 11). Ile starvation in *P. falciparum* was previously reported to induce a state of hibernation through the arrest of protein synthesis from which parasites could be recovered (*31*). We found that expression of PfPelo and Hsp70 was upregulated in starved cells (Fig. 4A and B) with a different localization pattern of the PfPelo protein in polysome profiles from the starved and controlled parasites (Supplementary Figure 11). Interestingly, while PfPelo was recruited to polysome fractions in samples obtained from Ile starved *P. falciparum* cells, we did not observe targeted degradation, but somewhat stabilization of Ile-rich transcripts: such as palmitoyltransferase DHHC8 (PF3D7_1321400), as compared to non-starved control samples. These results indicate that NGD components are being actively recruited to the stalled ribosomes; however, this process is independent of endonucleolytic cleavage of stalled mRNAs observed in the other organisms.

**Figure 4.**
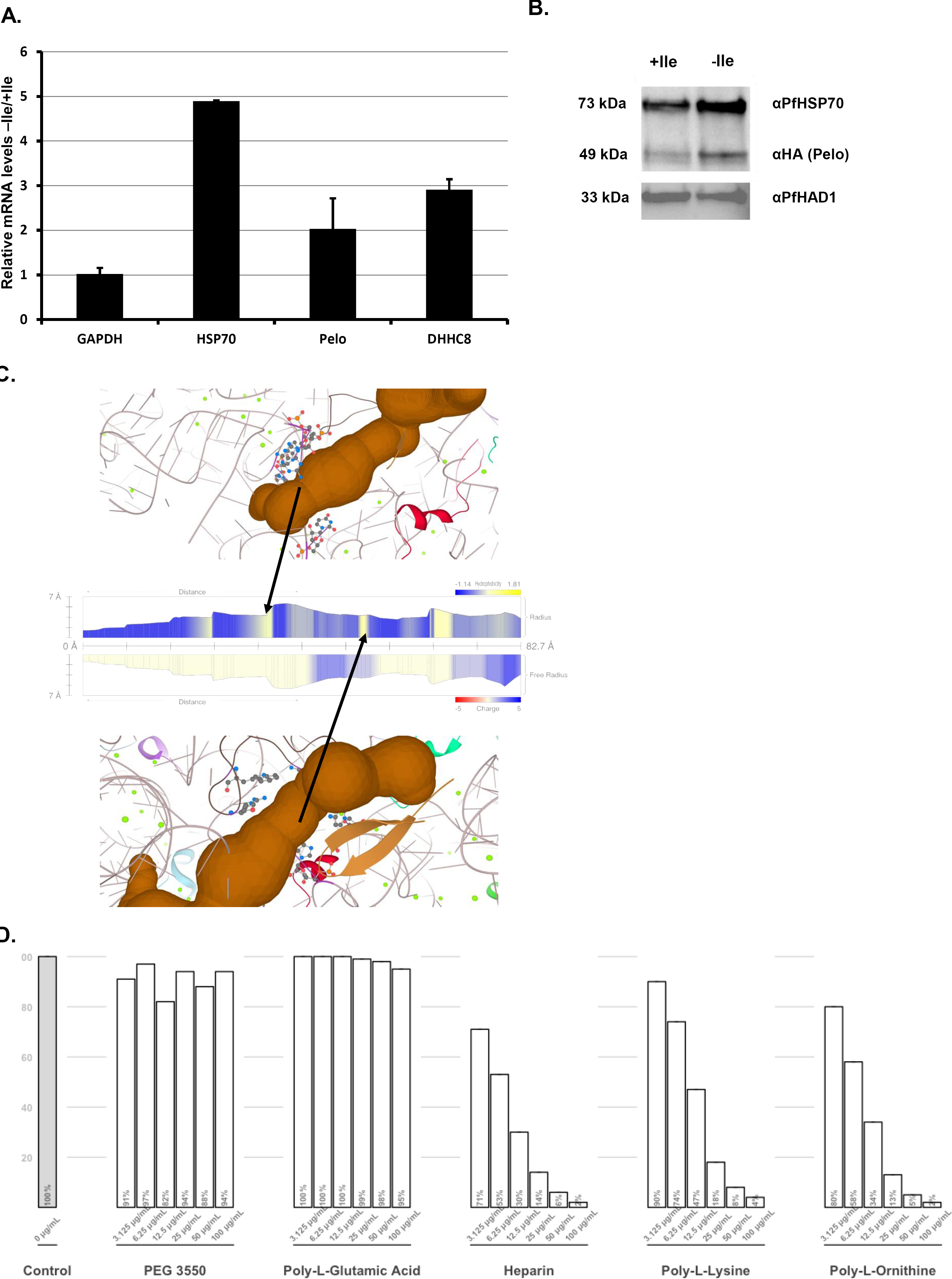
**A.** Relative mRNA enrichment for P. falciparum GAPDH (PF3D7_1462800), HSP70 (PF3D7_0818900), Pelo (PF3D7_0722100) and palmitoyltransferase (DHHC8; PF3D7_1321400) transcripts after 48 hours isoleucine (Ile) starvation of the *P. falciparum* cells. DHHC8 transcript codes for five consecutive Ile-residues. Levels of each transcript are normalized to total ribosome levels and represented as a ratio of transcript levels under Ile starvation (-Ile) over the control conditions (+Ile). **B.** Increased levels of P. falciparum HSP70 and Pelo proteins after 48 hours of Ile starvation. Western blot analysis of Ile starved (-Ile), and control (+Ile) sample are normalized to PfHAD1 (PF3D7_1033400) levels. HA-tagged *P. falciparum* Pelo protein was CRISPR/Cas9 engineered and detected using mouse HA-antibody. The molecular weight of each protein is indicated **C.** Polypeptide exit channel from Plasmodium ribosome (PDB: 3j79) has relatively short fragments (up to 3 Angstroms) of the relatively hydrophobic lining of the tunnel. Two major patches observed in other organisms are rRNA flanked (upper panel, interacting molecules have atoms shown) and L22 and L4 flanked (lower panel). **D.** *P. falciparum* growth assay in the presence of different compounds and amino acid polymers. Data represent endpoint parasitemia after 72-hour treatment with the indicated compound. Results are normalized to non-treated cultures (control) and presented as a percentage of control samples. Average value from three biological replicates is reported with standard deviation.

Recently, *P. falciparum* ribosomes were also shown to have different structural and dynamic features that distinguish them from other organisms; with the probable absence of RACK1 being one of the most prominent features regarding the translation of polyA tracks and poly-lysine repeats (*32, 33*) (Supplementary Fig. 12). Recent reports from yeast cells indicate the requirement of RACK1 proteins for endonucleolytic cleavage of stalled mRNAs (*34*). Additionally, deletion of RACK1 in human and yeast cells was also shown to increase production of proteins with polybasic peptides, however at the cost of increased frameshifting (*15, 35, 36*). To further investigate whether additional differences in the ribosome structure contributed to *P. falciparum* adaptation to long polyA tracks and poly-lysine repeats, we have compared ribosome exit channels between human, yeast, extremophile archaea and Plasmodium (Figure 4C and Supplementary Figures 13-15). Interestingly, two conserved patches of hydrophobic residues lining the exit channel (one being flanked by rRNA, while the other by ribosomal proteins L4 and L22) present in all four structures are almost barely noticeable in case of Plasmodium (Figure 4C). Given how unfavorable interactions are between clusters of positive charges and hydrophobic environments (*37*), this feature represents an interesting optimization of *P. falciparum* ribosomes that enables them to translate long poly-lysine repeats.

Conserved evolutionary adaptations result in traits that are beneficial for organisms. The increased AT-richness of *P. falciparum,* as such, could be a result of selective pressure on biosynthetic pathways (AT vs. GC biosynthesis) and the oxidative intraerythrocytic environment (*38*). Both of these environmental or cellular features would drive the genome towards higher AT content and, ultimately, increased polyA track length. However, we wondered whether benefits of *Plasmodium*’s increase in poly-lysine repeats might have different etiology. Our gene ontology analyses of polyA track and polylysine genes in *P. falciparum* suggested cellular and pathological adhesion proteins as one of the enriched gene groups between all exported proteins from *P. falciparum* (Supplementary Tables 1 and 2, respectively). Calculated isoelectric points for both groups are high, on average, and do not differ significantly from each other or *P. falciparum* proteins (Supplementary Fig. 16). This feature seems to be conserved to exported proteins of all *Plasmodium* species and certain parasitic species of *Microsporidia* (Supplementary Fig. 17). The lack of *P. falciparum* taxis during invasion of host cells, alterations in cytoadhesive properties of infected erythrocytes (*39*) and previously shown invasion-blocking effects of heparin (*40*) prompted us to test whether poly-lysine repeats in *P. falciparum* membrane proteins contribute to the overall efficiency of parasite invasion of host erythrocytes. Incubation of parasites with equal concentrations of short poly-(L) lysine or poly-(L) ornithine repeats in the media for 72 hours resulted in the efficient reduction of parasite growth for both poly-basic compounds (Fig. 4D). The range of the effect for both polymers was similar to previously reported inhibitory effects of heparin on *P. falciparum* invasion of human erythrocytes (*40*), which served as an experimental control. Addition of poly-glutamate or poly-ethylene glycol did not affect the growth of parasites; arguing that poly-basic polymers and heparin are explicitly inhibiting parasite growth. These data demonstrate that addition of poly-lysine exogenously to human erythrocyte cultures can prevent parasite growth in erythrocytes; possibly through competition with endogenously expressed poly-lysine and other polybasic proteins.

Our data indicate that the *P. falciparum* translational machinery permits the translation of polyA tracks without mRNA degradation, protein attenuation, nor activation of translational surveillance pathways. It is apparent that the accommodation of polyA tracks and poly-lysine repeats in translation required multiple adaptations to be vital, and typically highly conserved components of the translational and mRNA surveillance pathways; demonstrated here. While adaptations to polyA tracks have shaped translational accuracy and efficiency of *P. falciparum* ribosomes, it is indicative that ribosome exit channel differences might be adaptations to poly-lysine and other poly-basic sequences in *Plasmodium*, based on data from other organisms (*9, 41, 42*). Changes in the mRNA surveillance cellular components and mechanism might have followed ribosomal adaptations. However, increase in polyA tracks in DNA and mRNA sequences could have been driven by different selective pressure than the establishment of poly-lysine repeats in proteins. Advantages of parasites to synthesize proteins with poly-lysine repeats could be solely driven by benefits in the adhesion and invasion of host cells. Enrichment of proteins with poly-lysine or poly-basic repeats at the *P. falciparum* merozoite membrane is necessary to make initial adhesion of inert parasite cells to the erythrocyte membrane and increase the efficiency of invasion (*43, 44*). Also, it was suggested that an expansion of lysine-rich repeats in *Plasmodium* is associated with increased protein targeting to the erythrocyte periphery as seen with other poly-basic peptides (*45-47*). However, the conservation of both polyA tracks and poly-lysine repeats within *Plasmodium* species argue against the possibility that these sequences are just signals for plasma membrane localization and cell-surface retention. It is also possible that poly-lysine repeats in proteins expressed on the surface of infected erythrocytes contribute to previously observed sequestration of infected erythrocytes in particular organs of the human (*44*). Our data on the suppression of parasitic growth using exogenous poly-basic peptides indicate that such peptides could potentially be used as new antimalarial drugs. Further insights into differences between components of translational machinery and mRNA surveillance pathways present in *P. falciparum* and host organisms, will provide additional new drug targets against malaria.

## Acknowledgements

We thank members of Daniel Goldberg’s and Sergej Djuranovic’s lab for helpful comments. This work is supported by NIH R01 GM112824 to SD, NIH R00 R00GM112877 to WM, NSF MCB 1412336 to DLC, and NIH T32 GM: 007067 to JE. POB and JAJF are supported by the Washington University Center for Cellular Imaging (WUCCI) which is funded by Washington University School of Medicine, The Children’s Discovery Institute of Washington University and St. Louis Children’s Hospital (CDI-CORE-2015-505) and the Foundation for Barnes-Jewish Hospital (3770). SPD, JE and SD hold US Provisional Patent #62/696,868 titled “Antimalarial compositions and methods of use thereof.” The authors declare that they have no competing interests.

Author’s contribution: SPD and JE designed and conducted experiments, analyzed and interpreted biochemical data and drafted the article; RJA, WM, JE and PS conducted, analyzed and interpreted bioinformatics data; SPD, POB, and JAJF conducted, analyzed and interpreted microscopy data; JJC and DLC, conducted experiments, analyzed and interpreted biochemical data from *T. thermophila*; SD, SPD, PS, and JE are responsible for conception and design of the study. All the authors contributed to the revision of the article.

